# Competition for cysteine acylation by C16:0 and C18:0 derived lipids is a global phenomenon in the proteome

**DOI:** 10.1101/2022.05.27.492351

**Authors:** Hana Nůsková, Fabiola Garcia-Cortizo, Lena Sophie Schwenker, Marcel Tiebe, Martin Schneider, Dominic Helm, Carissa Reid, Annette Kopp-Schneider, Aubry K. Miller, Aurelio A. Teleman

## Abstract

*S*-acylation is a reversible posttranslational protein modification consisting of attachment of a fatty acid to a cysteine via a thioester bond. Research over the last few years has shown that a variety of different fatty acids, such as C16:0, C18:0 or C18:1, are used in cells to S-acylate proteins. We recently showed that GNAI proteins can be acylated on a single residue, Cys3, with either C16:0 or C18:1 and that the relative proportion of acylation with these fatty acids depends on the level of the respective fatty acid in the cell’s environment. This has functional consequences for GNAI proteins, with the identity of the acylating fatty acid affecting the subcellular localization of GNAIs. Unclear is whether this competitive acylation is specific to GNAI proteins or a more general phenomenon in the proteome. We perform here a proteome screen to identify proteins acylated with different fatty acids. We identify 218 proteins acylated with C16:0 and 308 proteins acylated with C18-lipids, thereby uncovering novel targets of acylation. We find that most proteins that can be acylated by palmitic acid (C16:0) can also be acylated with C18-fatty acids. For proteins with more than one acylation site, we find that this competitive acylation occurs on each individual cysteine residue. This raises the possibility that the function of many different proteins can be regulated by the lipid environment via differential *S*-acylation.

## Introduction

Metabolites can directly regulate signaling pathways via posttranslational modification of proteins (Figlia et al., 2020). Examples include protein acetylation, *O*-GlcNAcylation or palmitoylation; each is the covalent attachment of a metabolite to a protein, which thereby modifies the protein’s function. We focus here on *S*-acylation, the reversible attachment of fatty acids to cysteines via a thioester bond. This modification is mostly present on membrane-associated and integral membrane proteins, where it increases a protein’s affinity for lipid compartments or specific membrane microdomains, consequently affecting its activity, stability, and function (Chamberlain and Shipston, 2015; Chen et al., 2018). Hence, *S*-acylation plays a role in both physiological processes such as cell differentiation (Zhang and Hang, 2017), cell death (Fritsch et al., 2021), apoptosis and phagocytosis (Dixon et al., 2021), as well as pathological processes such as neurological disorders (Zareba-Koziol et al., 2018), cardiac dysfunction (Essandoh et al., 2020) and cancer (Ko and Dixon, 2018).

The creation of a thioester bond between a fatty acid and a cysteine residue of a protein is catalyzed mainly by the family of zinc finger DHHC domain-containing protein acyl transferases (ZDHHC-PATs), with 23 members in humans (Rana et al., 2019). The reverse process that leads to the removal of a fatty acid is catalyzed by thioesterases, which include palmitoyl protein thioesterases (PPTs), acyl protein thioesterases (APTs) and α/β hydrolase domain-containing 17 proteins (ABHD17s) (Won et al., 2018). Acylated proteins therefore undergo continuous cycles of acylation and deacylation due to the opposing activities of acyl transferases and thioesterases. Deacylation occurs everywhere in the cell whereas acylation occurs mainly at the Golgi (Rocks et al., 2010).

A variety of different fatty acids can be attached to proteins via *S*-acylation. Since palmitate is the most common fatty acid on proteins (Muszbek et al., 1999; Schulte-Zweckel et al., 2019) *S*-acylation was originally referred to as ‘palmitoylation’.

Nonetheless other saturated and unsaturated fatty acids of varying carbon chain lengths can modify proteins, including palmitoleate (C16:1) (Schulte-Zweckel et al., 2019; Zheng et al., 2016), stearate (C18:0) (Fujimoto et al., 1993; Liang et al., 2002), oleate (C18:1) (Montigny et al., 2014), and arachidonate (C20:4) (Hallak et al., 1994; Liang et al., 2001). Metabolic labeling with fatty acid analogs containing a “clickable” functional group (Charron et al., 2009; Hang et al., 2007) has enabled the identification of which fatty acids modify which proteins (Suazo et al., 2021).

One question of interest is whether one protein species (e.g. KRAS) is modified by one type of fatty acid (e.g. palmitic acid) or whether one protein species can be alternatively acylated with different fatty acids. If differential acylation takes place, does this lead to different functional consequences depending on the identity of the fatty acid? We recently showed that GNAI proteins can be differentially acylated by either palmitic acid or oleic acid on one amino acid, Cys3 (Nuskova et al., 2021). The identity of the fatty acid acylating GNAI proteins has functional consequences, since acylation with palmitic acid, but not with oleic acid, shifts GNAI proteins into detergent-resistant fractions of the cell membrane where they potentiate EGFR signaling. We found that the relative stoichiometry of acylation with either palmitic acid or oleic acid on Cys3 depends on the relative abundance of the two fatty acids in the medium, thereby linking cellular lipid availability to a regulatory effect on GNAI function and EGFR signaling. Although we showed that this competitive acylation with different fatty acids occurs for one protein species (GNAI proteins), the question remains whether this is a general phenomenon also for other proteins. If this is the case, this would open up the possibility that the function of many different proteins can be regulated by differential acylation.

We report here a proteomic analysis of human *S*-acylated proteins focusing on differential acylation with palmitate versus stearate. Via metabolic labeling with azido fatty acid analogs of palmitate and stearate, we identify and validate already known *S*-acylated proteins as well as new targets of *S*-acylation. Analysis of differential *S*-acylation reveals that most *S*-acylated proteins can be modified by either fatty acid on single cysteine residues, indicating that competitive *S*-acylation with different fatty acids is a general phenomenon proteome-wide. This raises the possibility that the function of many proteins may be regulated in this manner.

## Results

### Most proteins S-acylated with C15-az can also be acylated with C17-az

To study protein *S*-acylation by fatty acids of two different lengths, namely palmitic acid (C16:0) and stearic acid (C18:0), we employed an established metabolic labeling assay that uses fatty acid analogs with an azide group at the methyl end, which makes them suitable for click chemistry reactions (**Figure 1**). Cells use these fatty acid analogs to acylate endogenous proteins *in vivo*. The proteins which have been *S*-acylated with a specific fatty acid can subsequently be purified and analyzed either by mass spectrometry or immunoblotting (Suazo et al., 2021). We previously showed that C17:0-azide (C17-az) is functionally equivalent to C18:0 (Nuskova et al., 2021; Senyilmaz et al., 2015). The C16:0 analog C15:0-azide (C15-az) has been extensively used to identify palmitoylated proteins (Charron et al., 2009; Hang et al., 2007). To metabolically label the proteome, we incubated cells with either BSA-conjugated azido fatty acids for 3 h, or with BSA only as a negative control for non-specific background. This leads to labeling of both *de novo* synthesized proteins as well as proteins that undergo deacylation/reacylation cycles. For instance, RAS has a protein half-life of 24 h but an acylation half-life <30 min (Ahearn et al., 2018; Lin and Conibear, 2015), GNAI proteins have a half-life >12 h but an acylation half-life of 3 h (Nuskova et al., 2021), and additional examples of proteins with rapid acylation turnover have been reported (Degtyarev et al., 1993; El-Husseini Ael et al., 2002; Zhang et al., 2010). We previously found that within this 3-hour timeframe, C17:0-azide or C18:0 can be unsaturated in the cell, yielding proteins carrying acylation with C17:1-azide or C18:1, respectively (Nuskova et al., 2021). In contrast, C15:0-azide and C16:0 remain predominantly saturated. For simplicity we use here the term “acylated with C17-az” or “with C18” to refer to proteins acylated with either the saturated or desaturated forms of the fatty acids. After metabolic labeling, we lysed the cells, captured the proteins labeled with azido fatty acids on agarose beads carrying alkyne groups via copper-catalyzed azide-alkyne cycloaddition (CuAAC), and analyzed them by label-free quantitative (LFQ) mass spectrometry (**Figure 1A**). This revealed 218 proteins enriched in the C15-az labeled samples and 308 proteins enriched in the C17-az labeled samples compared to the negative control (**Figure 1B-C**, **Supplementary Data 1**). These numbers were obtained by combining the proteins that were present exclusively in the C15-az or C17-az-labeled samples and not detected at all in any of the negative control replicates (“clean” proteins) together with the ones that were enriched compared to the negative control with an adjusted p-value<0.05 based on the biological replicates (**Figure 1C**). The majority of proteins that were labeled with C15-az were also labeled with the longer C17-az (**Figure 1D**). Roughly 80% of the proteins we identified as being acylated with either C15-az or C17-az were previously reported to be acylated according to the acylation database SwissPalm (Blanc et al., 2019) (**Figure 1D**). This percentage was similar for “clean” proteins and for the ones with p<0.05, suggesting that both categories of acylation targets are equally enriched for real acylation targets.

**Figure 1:**
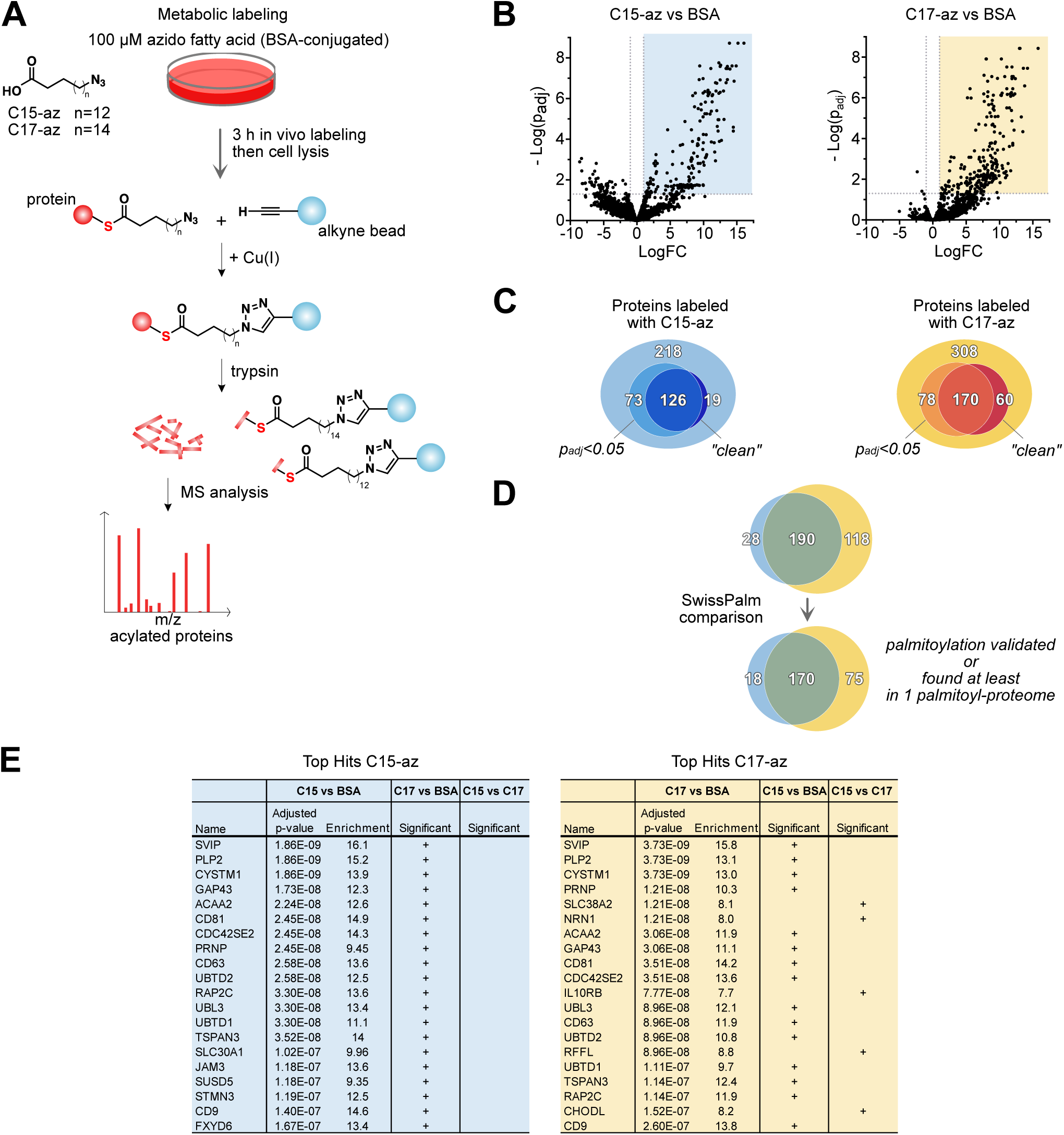
Detection of *S*-acylated proteins via metabolic labeling with azido fatty acids. A. Schematic diagram representing metabolic labeling with azido fatty acids followed by a pulldown of labeled proteins based on click chemistry. HEK293E cells were incubated with BSA-conjugated C15-az or C17-az (final concentration 100 μM) or BSA alone as a negative control for 3 h. Cells were lysed and incubated for 16–20 h with alkyne agarose under conditions of CuAAC-based click reaction. On-bead reduction, alkylation of proteins, and trypsin digest were performed followed by MS analysis of eluted peptides. B. Volcano plots showing proteins detected in C15-az (n=4) and C17-az (n=4) labeled samples in comparison to the negative control (BSA; n=5). The upper right quadrant in both graphs depicts proteins significantly enriched in C15-az or C17-az labeled samples (adjusted p-value<0.05 according to the statistical analysis of 2196 filtered proteins described in the Methods section). C. Venn diagrams showing the overlap of statistically enriched proteins in C15-az and C17-az labeled samples with proteins that were exclusively detected in the respective labeled samples and completely missing in the negative controls (“clean” proteins). The merged protein lists were then compared with the data contained in the SwissPalm database. D. Top 20 significantly enriched proteins in C15-az (left) and C17-az (right) labeled samples. In addition to the adjusted p-value and fold change, shown is whether the protein is also significantly enriched in the samples labeled with the other azido fatty acid and if there is a significant difference between these two labeling conditions.

The top hits in our proteomic screen include novel putative targets of *S*-acylation in addition to proteins known to be *S*-acylated (**Figure 1E**, **Supplementary Table 1**). We found small GTPases – subunits of trimeric G-proteins (GNAI1, GNAI2, GNAI3, GNAQ, GNAS, GNA11, GNA13) and Rho-family of GTPases (RHOB, RHOU, RHOQ). We also identified several SNARE proteins that play a key role in vesicle trafficking (Han et al., 2017), such as syntaxins (STX4, STX5, STX6, STX7, STX8, STX10, STX12), vesicle associated proteins (VAMP2, VAMP3, VAMP4, VAMP7) and others (SNAP23, SNAP25, TRAPPC3). In addition, our list of putative *S*-acylated proteins includes plasma membrane transporters (SLC1A5, SLC7A1, SLC7A2, SLC19A1, SLC30A1, SLC38A2, SLC39A6, SLC44A1) and also some death receptors belonging to the tumor necrosis factor receptor superfamily (TNFRSF) (Fas/CD95, DR4/TRAIL-R1, DR5/TRAIL-R2, DcR2/TRAIL-R4) some of which were previously reported (Fritsch et al., 2021). Our screen also indicates that a component of the insulin signaling pathway, IRS4, can be directly modified by fatty acids. IRS4 acylation has not been reported previously most likely due to its limited tissue-specific expression. HEK293 cells are one of the few cell lines that express this protein. Last, we found some proteins involved in the acylation/deacylation machinery including members of the ZDHHC family of acyl transferases (ZDHHC3, ZDHHC18, ZDHHC20), and two thioesterases, PPT1 and ABHD17B (FAM108B1), whose *S*-acylation has been validated in other studies (Martin and Cravatt, 2009; Segal-Salto et al., 2016).

Taken together, our analysis of metabolically labeled proteins identified both proteins already known to be *S*-acylated as well as many new putatively *S*-acylated proteins, which we study further below. Our results indicate that the majority of *S*-acylated proteins can be modified with both C15-az and C17-az.

### Validation of putatively S-acylated proteins by APE assay

We next validated the putative *S*-acylation of proteins we identified in our proteomic screen using an independent method. For this, we used the acyl-PEG exchange (APE) assay, which entails lysing cells, blocking free cysteines with *N*-ethylmaleimide (NEM), and subsequently cleaving the acyl-thioester bonds with hydroxylamine. After removal of fatty acids, the newly generated free cysteines can be specifically labeled with thiol-reactive compounds linked to mPEG, which causes a mass-shift on an SDS-PAGE gel (Drisdel and Green, 2004; Percher et al., 2016; Roth et al., 2006; Wan et al., 2007). Since the elution step uses hydroxylamine, which specifically cleaves thioester bonds, this assay distinguishes *S*-acylation from other types of acylation, such as *N*-acylation, which do not take place via a thioester bond. We used the APE assay because it can also reveal the number of *S*-acylated sites if the antibody is of a good quality (Percher et al., 2016).

Using this assay, we validated the *S*-acylation of multiple novel *S*-acylation targets (**Figure 2A-P**). This includes proteins that were identified in both the C15-az and C17-az screens, as well as proteins whose enrichment was statistically significant only for one azido fatty acid (C15-az: IRS4 and SLC1A5; C17-az: CANX and RFFL) (**Figure 2A-P**). Although the antibody seemed to work well, we did not observe *S*-acylation of STX5 (**Figure 2Q**), which suggests that it might be a false positive or a different kind of acylation such as *N*-acylation. We also tested some proteins that were enriched in our proteomic screen but missed our significance cut-off, and found that they are also *S*-acylated, such as CYB5B and FASN (**Figure 2R-S**). Although this suggests that our statistical cut-offs may be overly stringent, we prefer to have a dataset that is smaller but has few false positives. Indeed, using the APE assay we validated approximately 80% of all the proteins that we tested, including multiple new *S*-acylation targets.

**Figure 2:**
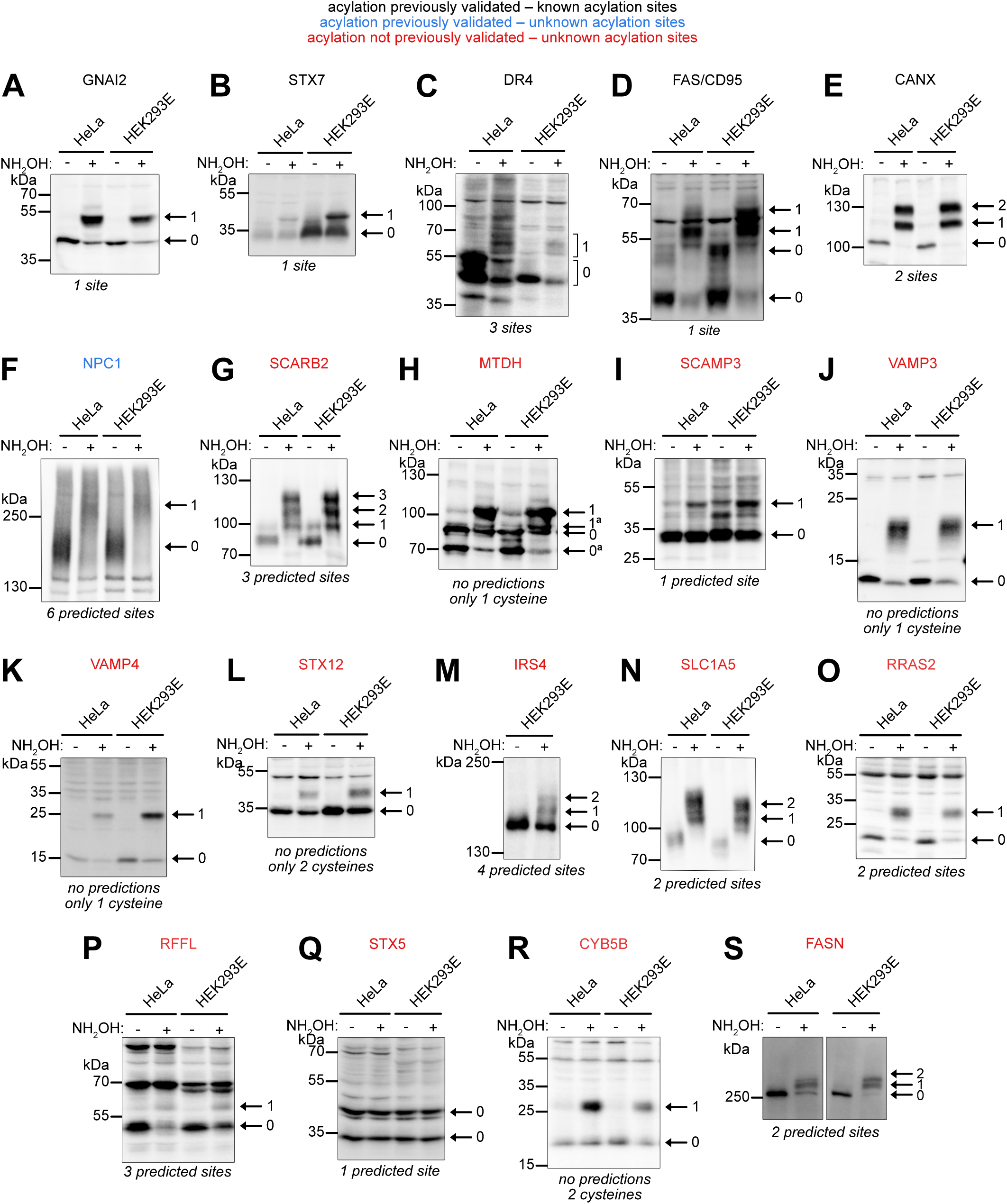
Validation of putative *S*-acylated proteins by acyl-PEG exchange (APE) assay. (**A-S**) Validation of *S*-acylation of proteins enriched in C15-az and C17-az labeled samples using the APE assay. Proteins whose *S*-acylation had been previously validated in other studies were included as positive controls of the technique (labeled in black – known acylation sites, and in blue – unknown acylation sites). Putative *S*-acylated proteins that were not validated before are labeled in red. The *S*-acylated isoforms are indicated on the right side of each Western blot (0 – non-acylated protein, 1 – protein with 1 *S*-acylated cysteine etc.), another isoform of a different molecular weight is also indicated if present (panel h). Under each Western blot, the information on the number of known or predicted *S*-acylation sites is given for already validated and putative *S*-acylated proteins, respectively (according to the SwissPalm). Representative Western blots of at least 3 biological replicates.

### Different fatty acids compete to acylate single cysteine residues

We next asked whether competitive acylation of a protein with different fatty acid species is a general phenomenon. We noticed that some proteins that can be modified by different fatty acids (e.g. STX7, CYB5B, VAMP3, VAMP4) (**Suppl. Table 1**), have only one acylation site, as can be seen by the fact that they only give a single up-shifted band in the APE assay (**Figures 2B, K, L** **and** **R**). Indeed, VAMP3 and VAMP4 only have one cysteine in the whole protein. Hence, for these proteins the two different fatty acids must compete for acylating this one site. However, there are many proteins with multiple acylation sites. In these cases, it is possible that one fatty acid (e.g. C15-az) acylates one site on the protein and a different fatty acid (C17-az) acylates another site. Alternatively, the different fatty acids might compete with each other to acylate each site on the protein. To distinguish these two possibilities, we chose two model proteins whose post-translational modifications are well known – LAMTOR1 and HRAS – and tested whether there is heterogenous *S*-acylation on each site, as described below.

LAMTOR1 (also known as p18 or c11orf59) plays a key role in mTORC1 signaling. As a member of the Ragulator complex, LAMTOR1 takes part in recruitment of mTOR to lysosomes upon amino acid stimulation (Sancak et al., 2010). Since LAMTOR1 is the only member that is modified by lipids, it provides an essential anchor for localization of the whole Ragulator complex and mTORC1 to lysosomes (Martin and Cravatt, 2009; Nada et al., 2009). There are only two cysteine residues in LAMTOR1, both at the N-terminus, located directly next to each other (**Figure 3A**). Their *S*-acylation follows a proteolytic cleavage of the N-terminal methionine and subsequent *N*-myristoylation of the adjacent glycine (Sanders et al., 2019). By APE assay, we confirmed that LAMTOR1 is acylated on two cysteine residues (**Figure 3B**), and that the two cysteines are Cys3 and Cys4 since the double-mutant loses all acylation (**Figure 3C**). Acylation of each cysteine is not dependent on the other, since each single cysteine mutant retains acylation on the other site (**Figure 3C**). Metabolic labeling with azido fatty acids of the single mutants (C3S or C4S) revealed that each site can be acylated by either C15-az or C17-az (**Figure 3D**). We used GNAI2 and TRF1 as positive controls for the *in vivo* labeling and pulldown. LAMTOR1 labeling is even more efficient in the case of the single cysteine mutants compared to wild-type LAMTOR1, which could be due to increased turnover of *S*-acylation on LAMTOR1 when only one cysteine is modified.

**Figure 3:**
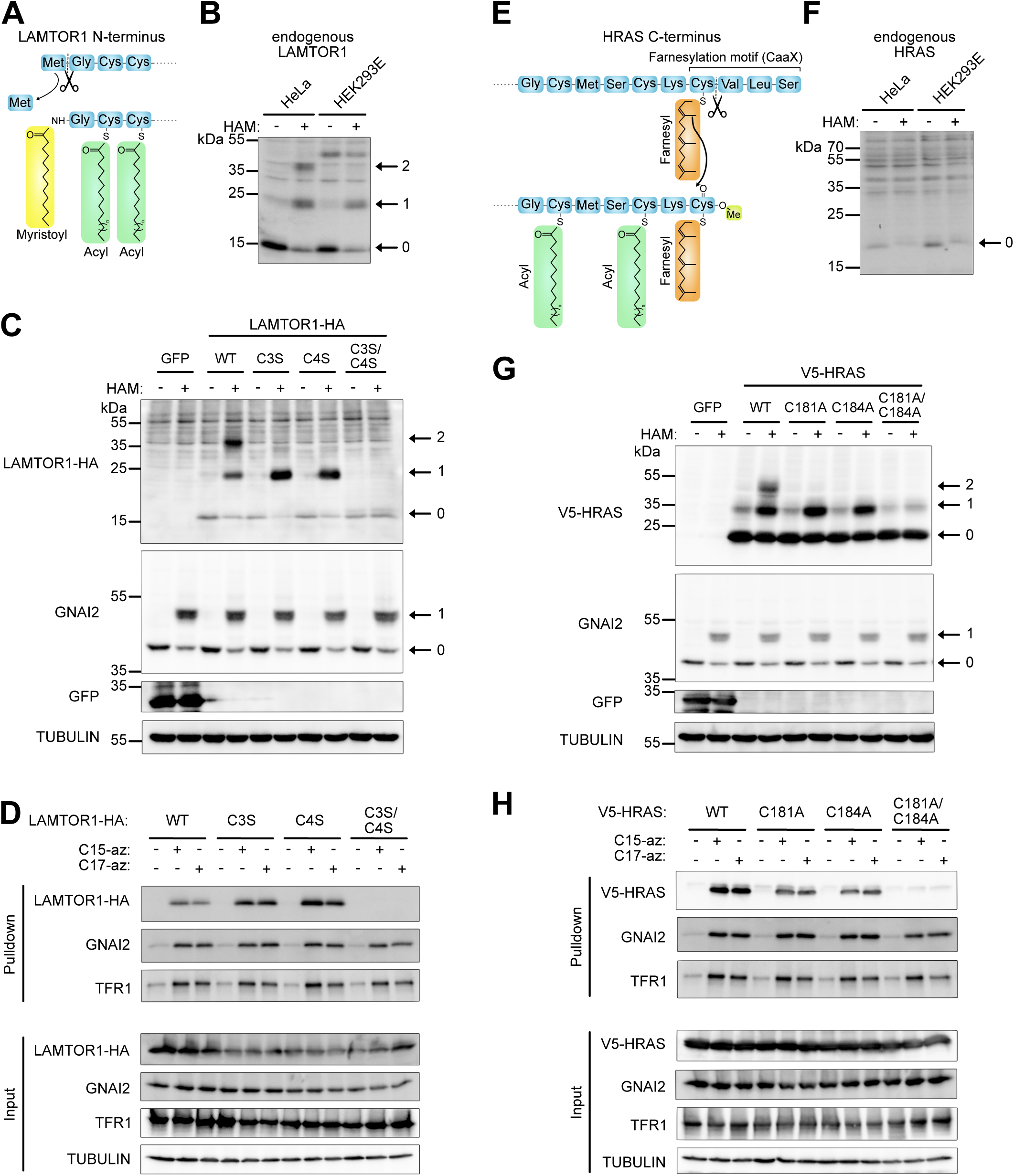
LAMTOR1 and HRAS as examples of differential *S*-acylation on multiple acylation sites. A. A scheme illustrating the N-terminal sequence of LAMTOR1 with its lipid modifications. B. APE assay detects acylation sites on the endogenous LAMTOR1 in HeLa and HEK293E cells. A representative image of 3 biological replicates. C. APE assay detects that both Cys3 and Cys4 are acylated in LAMTOR1-HA. HeLa cells were transiently transfected with overexpression constructs and lysed for APE assay 24 h post transfection. Wild-type LAMTOR1 was compared to acylation mutants upon detection with anti-HA antibody (single mutants C3S, C4S and the double mutant C3S/C4S). The endogenous GNAI2 with its single acylation is shown as a positive control of the assay and tubulin as a loading control. A representative of 3 independent experiments. D. Both Cys3 and Cys4 are acylated by either C15-az or C17-az in LAMTOR-HA. Metabolic labeling with azido fatty acids (conditions as in Figure 1A) in HeLa cells was performed 24 h after transfection. The purified labeled proteins were eluted for a Western blot and the wild-type LAMTOR1 was compared to the acylation mutants (detected with anti-HA antibody). The efficiency of GNAI2 and TFR1 labeling serves as a positive control of the assay. A representative of 2 independent experiments. E. A scheme illustrating the C-terminal sequence of HRAS with its lipid modifications. F. APE assay fails to show acylation sites on the endogenous HRAS in HeLa and HEK293E cells, likely due to a faint antibody signal. A representative image of 3 biological replicates. G. Both Cys181 and Cys184 are acylated in V5-HRAS, detected via APE assay. Transiently transfected HeLa cells were lysed for APE assay 24 h post transfection. Wild-type HRAS was compared to acylation mutants upon detection with anti-V5 antibody (single mutants C181A, C184A and the double mutant C181A/C184A). The endogenous GNAI2 with its single acylation is shown as a positive control of the assay and tubulin as a loading control. A representative of 3 independent experiments. H. Both Cys181 and Cys184 are acylated by both C15-az and C17-az in V5-HRAS. Metabolic labeling with azido fatty acids (conditions as in Figure 1A) in HeLa cells was performed 24 h after transfection. The wild-type V5-HRAS was compared to the acylation mutants (detected with anti-V5 antibody). The efficiency of GNAI2 and TFR1 labeling serves as a positive control of the assay. A representative of 3 independent experiments.

We identified several members of the RAS superfamily of small monomeric guanosine triphosphatases (GTPases) in our pulldown of *S*-acylated proteins, namely NRAS, HRAS and a number of other RAS-related proteins (RRAS2, RAP2A, RAP2B, RAP2C, RAB6C) (**Suppl. Table 1**). RAS proteins comprise an important class of molecular switches that regulate cellular processes (Chen et al., 2021). The post-translational modifications on the C-terminus of RAS proteins are well known (Ahearn et al., 2018; Busquets-Hernandez and Triola, 2021). Briefly, after synthesis in the cytosol, their C-terminal CAAX motif is farnesylated on the cysteine residue leading to subsequent targeting to the ER membrane. At the ER, the AAX sequence is cleaved off and the carboxyl group of the new C-terminal cysteine is methylated. While the farnesyl residue already acts as a membrane anchor, before RAS proteins are transferred to the plasma membrane via the secretory pathway the membrane affinity can be increased by additional acylation at a proximal cysteine residue (Busquets-Hernandez and Triola, 2021). HRAS has two acylation sites, Cys181 and Cys184 (**Figure 3E**) (Hancock et al., 1989). Using the APE assay on endogenous HRAS, we could not detect any up-shifted bands, although we could observe a lower intensity of the band corresponding to non-acylated HRAS (arrow, **Figure 3F**), which underlines the requirement for a good antibody for APE assays. Nonetheless, we confirmed *S*-acylation of HRAS on Cys181 and Cys184 using overexpressed V5-tagged protein (**Figure 3G**). As for LAMTOR1, metabolic labeling with azido fatty acids using singly-mutated HRAS revealed that each cysteine can be modified by either C15-az or C17-az.

In conclusion, we find that different fatty acids compete to *S*-acylate single cysteine residues on proteins, and that if a protein has multiple *S*-acylation sites, this heterogenous *S*-acylation occurs on each individual acylation site. This suggests that heterogenous *S*-acylation is not a feature of a single acylation site but is instead a general phenomenon within the proteome. This raises the possibility that the function of many proteins could be regulated in this way.

### Some S-acylated proteins are more efficiently acylated with one particular fatty acid

Not all proteins were significantly enriched in both the C15-az and C17-az pulldowns (**Figure 1D**), suggesting that some proteins may be preferentially acylated with a specific fatty acid. To directly and quantitatively compare the labeling of proteins with different azido fatty acids, we employed stable isotope labeling with amino acids in cell culture (SILAC). Briefly, isotopically labeled lysine and arginine (“light”, “medium” and “heavy”) were incorporated into cellular proteins via prolonged cell culture prior to addition of C15-az, C17-az or unconjugated BSA as a negative control for 3 h (**Figure 4A**). Differentially labeled samples were then mixed prior to the click reaction, pulldown, and mass spectrometry. The isotope labeling was swapped in biological replicates of the experiment to exclude technical biases. We then statistically analyzed the data focusing on two aspects. First, proteins significantly enriched in C15-az and C17-az labeled samples relative to the negative control were determined (**Figure 4B, C**, **Supplementary Data 2**). The statistically enriched proteins (adjusted p-value<0.05) overlapped to a large extent with the proteins identified in the conventional screen described above. Second, for these proteins the relative stoichiometry of labeling with C15-az versus C17-az was compared and several were found to be preferentially labeled with one fatty acid compared to the other (**Figure 4D**). The magnitude of this effect, however, was mild and at most 2-fold, corresponding to 1/3 of the total protein being acylated with one fatty acid and 2/3 with the other, rather than half-half. More proteins were preferentially acylated with palmitate compared to C17-az: out of 32 proteins tested, 12 and 4 proteins were significantly enriched (unadjusted p-value<0.05) in C15-az and C17-az labeled samples, respectively. In sum, we observed that some proteins may be preferentially *S*-acylated with a certain fatty acid, but we did not find any that were exclusively labeled with one fatty acid but not the other.

**Figure 4:**
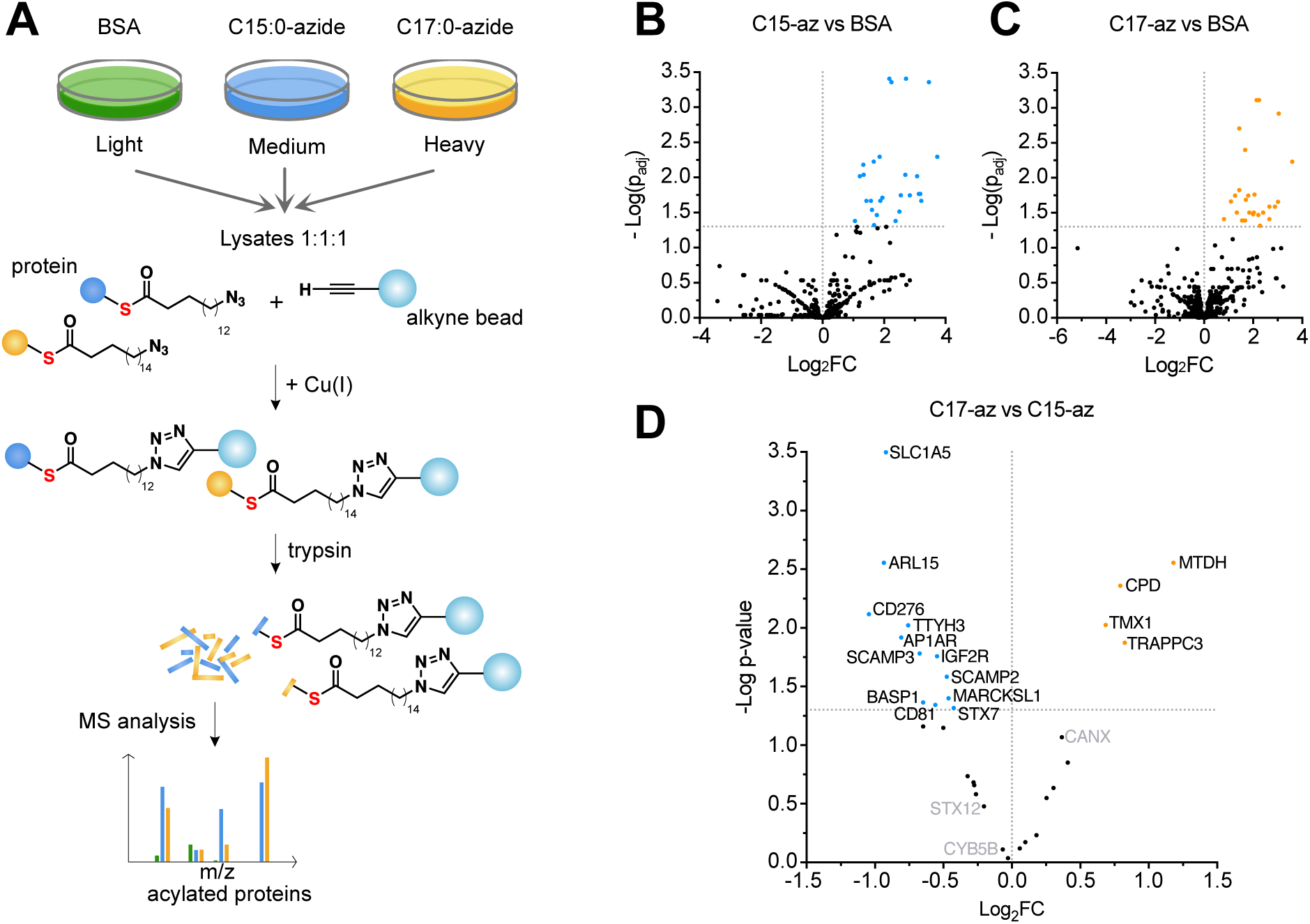
SILAC screen for preferential acylation of proteins with either C15-az or C17-az. A. Schematic representation of the metabolic labeling with azido fatty acids followed by a pulldown of labeled proteins based on click chemistry in combination with the SILAC approach. Prior to the metabolic labeling, HEK293E cells were cultured in medium containing isotopically labeled amino acids for an extended period of time (“light” – unlabeled amino acids; “medium” – ^2^H-labeled lysine and ^13^C-labeled arginine; “heavy” – ^13^C, ^15^N-labeled lysine and ^13^C, ^15^N-labeled arginine). To label *S*-acylated proteins, cells were incubated with BSA-conjugated C15-az or C17-az (final concentration 100 μM) or BSA alone as a negative control for 3 h. Cells were lysed, mixed equally and incubated for 16–20 h with alkyne agarose under conditions of CuAAC-based click reaction. In two other replicates, different conditions and labeling were swapped. On-bead reduction and alkylation of proteins and trypsin digest was performed before a subsequent MS analysis of eluted peptides. B-C. Volcano plots showing proteins detected in C15-az and C17-az labeled samples in comparison to the negative control (BSA) (n=3 for each SILAC-based ratio). The upper right quadrant in both graphs depicts proteins significantly enriched in C15-az or C17-az labeled samples (p<0.05). Blue (C15-az) and orange (C17-az) dots indicate proteins enriched with a high confidence (adjusted p-value<0.05; based on the statistical analysis described in the Methods section). D. A volcano plot showing proteins enriched in C15-az vs C17-az. The statistical analysis of C15-az/C17-az ratios was performed on proteins that were enriched in C15-az or C17-az labeled samples compared to the BSA control with p-value<0.05 (as shown in **B** and **C**).

To validate hits of the SILAC-based screen, we selected SCAMP3, SLC1A5 (ASCT2), and STX7 as proteins that are preferentially acylated with palmitate (C15-az), MTDH (LYRIC) and TMX1 as proteins that are preferentially acylated with C17-az, and CANX, CYB5B and STX12 as acylated proteins that did not display any significant differences between the two lipids (**Figure 4D**). From the conventional screen (**Figure 1D**), we selected RFFL, which demonstrated a preference for C17-az and whose *S*-acylation we confirmed by the APE assay (**Figure 2P**), and TFR1, since we repeatedly observed a higher efficiency of C15-az labeling in previous assays (**Figure 3**). Furthermore, we also tested proteins whose *S*-acylation we had already validated (**Figure 2**) but were either not detected in the SILAC assay, or did not display any significant preference: DR4 (TRAIL-R1), NPC1, VAMP3, and SCARB2 (LIMP2). To compensate for potential variability among samples due to technical reasons, we used GNAI2 for normalization since it is equally well labeled with both C15-az and C17-az (Nuskova et al., 2021). We treated cells with C15-az or C17-az to allow protein acylation *in vivo* and analyzed the pulldowns by immunoblotting. Using this approach, we could confirm all the expected differences in HeLa cells (**Figure 5A,B**). All proteins that showed increased acylation with C17-az preference in the SILAC screen, namely MTDH, TMX1, and RFFL, were more strongly acylated with C17-az also in this validation assay in HeLa cells (significant in the case of MTDH and TMX1) (yellow bars **Figure 5B**). All the proteins that were found more strongly acylated with C15-az compared to C17-az in the SILAC screen also behaved this way in the validation assay (blue bars **Figure 5B**). In addition, several proteins that were either not detected in the SILAC screen or showed no significant preference for acylation with one lipid, were nonetheless more strongly acylated with C15-azide compared to C17-azide (**Figure 5A-B**), suggesting that the SILAC screen may have missed some differentially acylated proteins. Nonetheless, the magnitude of the preference for one lipid or the other is mild in all cases, hardly exceeding 2-fold. The same tendencies were observed also in HEK293E cells (**Figure 5C,D**), indicating that the fatty acid preference of tested proteins is conserved in these two cell lines. In conclusion, we find that some proteins are preferentially acylated with C18 lipids, many proteins are more easily *S*-acylated with shorter palmitate, but the preference for one lipid or the other is mild in all cases.

**Figure 5:**
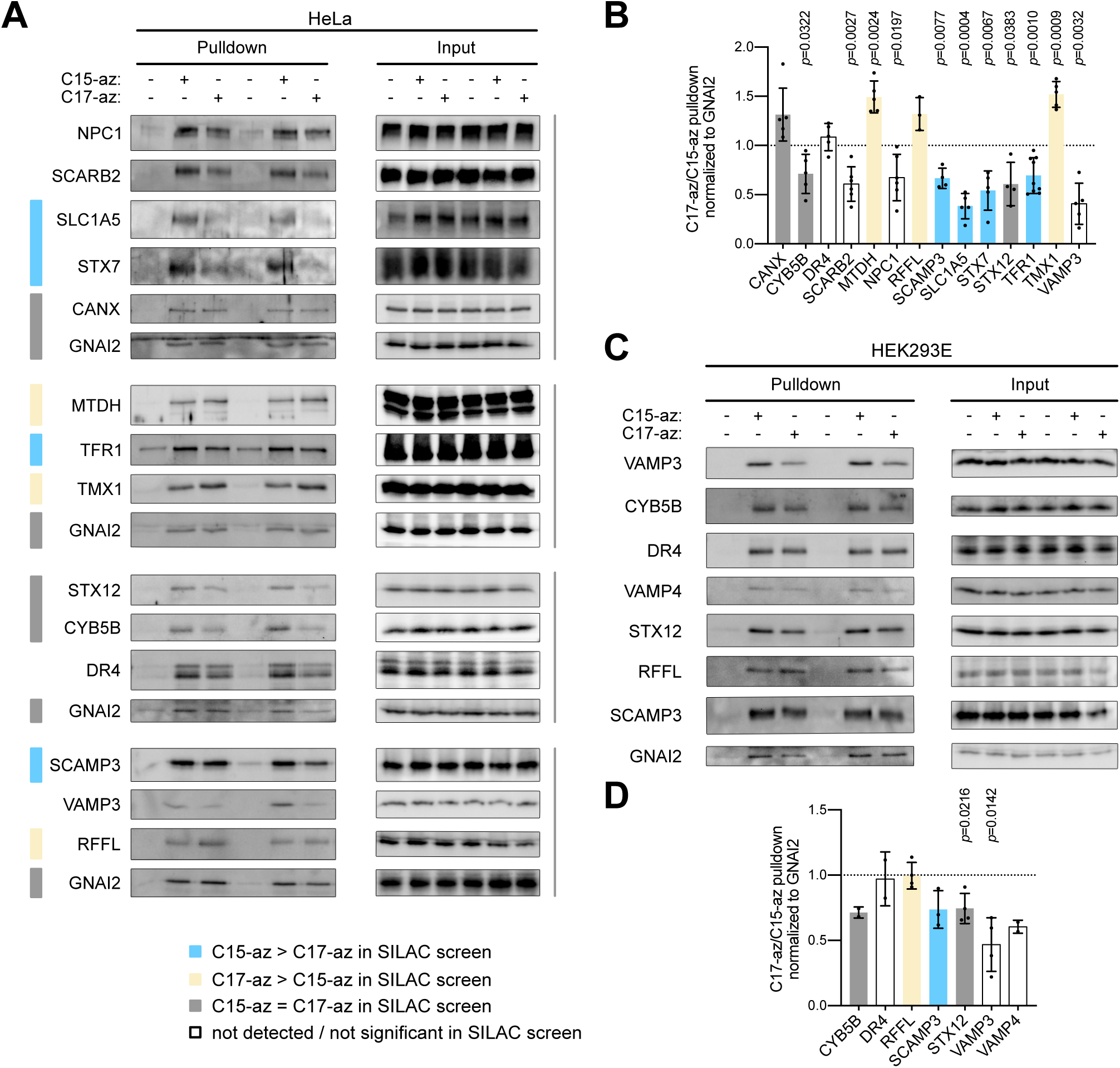
Validation of preferential *S*-acylation of proteins with one fatty acid. Metabolic labeling with azido fatty acids in HeLa (A-B) or HEK293E (C-D) cells was performed under the conditions described in Figure 1A. After the click-based pulldown on alkyne agarose, labeled proteins were eluted and probed on Western blots. The efficiency of GNAI2 labeling serves as a positive control of the assay. The intensity of bands was quantified, the values were background corrected and normalized to the respective GNAI2 band in both HeLa (**B**) and HEK293E (**D**) cells. The ratio of the efficiency of C17-az and C15-az labeling is shown for each tested protein. The significance was tested by one sample t-test (hypothetical mean 1.0), when significant, p-value is indicated, n=2–5 (weak antibodies could not be detected in some samples), mean±SD.

### Fatty acid preference is not a feature of an individual acylation site, but of the whole protein

For proteins that show stronger acylation with one fatty acid than another, this might be caused by one specific cysteine acylation site on the protein. Alternatively, all cysteines on the protein could show the same tendency, perhaps because of the protein’s subcellular localization. For proteins that only have a single acylation site this is a moot point. For example, STX7 (syntaxin 7), one of the endosomal SNARE proteins that mediates membrane fusion, is acylated on Cys239, which is adjacent to its transmembrane domain (He and Linder, 2009). Both the SILAC screen results (**Figure 4**) and the validation assay (**Figure 5**) indicate that this cysteine is more strongly acylated with palmitate rather than C18. Similarly, the palmitate preference was observed for singly-acylated CYB5B, STX12, VAMP3 and VAMP4 (**Figure 5**). To study a protein containing more than one acylation site, we selected TFR1 (transferrin receptor 1; also known as CD71), which is more strongly acylated with C15-az compared to C17-az (**Figure 5A,B**). TFR1 is an integral membrane glycoprotein with a single transmembrane domain. The TFR1-mediated endocytosis of diferric transferrin represents an important mechanism how cells uptake iron (Ciechanover et al., 1983). TFR1 is acylated on two sites, Cys62 and Cys67, which regulate its endocytosis (Alvarez et al., 1990) (**Figure 6A**). Using the APE assay we could detect acylation on both sites on the endogenous protein (**Figure 6B**). On overexpressed wild-type protein, however, dual acylation is rare (**Figure 6C**). Nonetheless, both sites can be acylated because both have to be mutated to abolish acylation completely (**Figure 6C**). Interestingly, both acylation sites of TFR1, Cys62 and Cys67, are more strongly acylated with C15-az compared to C17-az (**Figure 6D**), suggesting that the preference for one fatty acid is not a feature of an individual acylation site, but of the whole protein.

**Figure 6:**
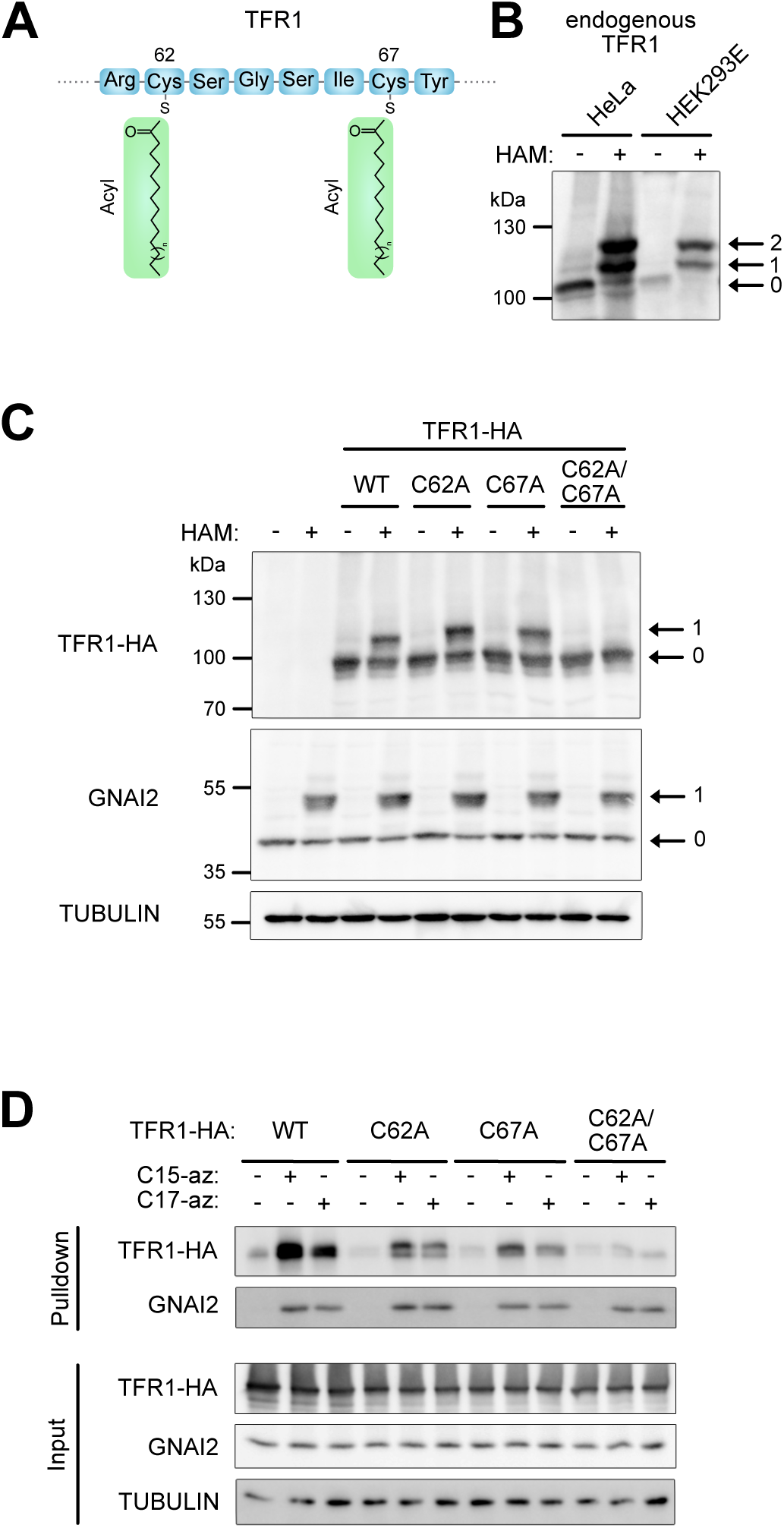
TFR1 as an example of preferential *S*-acylation on multiple acylation sites. A. Schematic diagram of amino acids 61-68 of TFR1, which contain lipid modifications. B. APE assay detects acylation sites on endogenous TFR1 in HeLa and HEK293E cells. C. Both Cys62 and Cys67 are acylated in TFR1-HA, detected via APE assay. Transiently transfected HeLa cells were lysed for APE assay 24 h post transfection. Wild-type TFR1 is compared to acylation mutants upon detection with anti-HA antibody (single mutants C62A, C67A and the double mutant C62A/C67A). The endogenous GNAI2 with its single acylation is shown as a positive control of the assay and tubulin as a loading control. A representative of 3 independent experiments. D. Both Cys62 and Cys67 are preferentially acylated with C15-az compared to C17-az. Metabolic labeling with azido fatty acids (conditions as in Figure 1A) in HeLa cells was performed 24 h after transfection. The wild-type TFR1-HA was compared to the acylation mutants (detected with anti-HA antibody). The efficiency of GNA2 labeling serves as a positive control of the assay. A representative of 2 independent experiments.

## Discussion

We previously showed that GNAI proteins can be acylated with either palmitic acid or oleic acid on a single cysteine residue, Cys3, and that the identity of that fatty acid affects the function of the GNAI protein (Nuskova et al., 2021). We aimed to study here whether such differential acylation of individual protein species with different fatty acids is a general phenomenon observed throughout the proteome. Our results indicate that indeed the majority of *S*-acylated proteins can be modified with either C15-az or C17-az. In cases where proteins have several *S*-acylation sites, we find that different fatty acids compete for *S*-acylation of each single cysteine residue. Therefore, heterogenous *S*-acylation is not a feature of a single acylation site but instead of the whole protein. We also observed that some *S*-acylated proteins are preferentially acylated with a certain fatty acid, often the shorter palmitate (C15-az), however this bias is small in magnitude. In sum, differential *S*-acylation of proteins may be a general mechanism how lipids regulate protein function in a cell.

Our results show that many proteins are more efficiently modified with C15-az than with the longer C17-az (**Figure 4**). Indeed, palmitate is the fatty acid that seems to be most frequently attached to proteins (Muszbek et al., 1999; Schulte-Zweckel et al., 2019). Other fatty acids, such as palmitoleate, stearate, or oleate, can be also utilized for protein *S*-acylation but the frequency of their use seems to be cell type dependent (Schulte-Zweckel et al., 2019). Although our knowledge on the regulation of the fatty acid selectivity of ZDHHC-PATs is still lacking, it is becoming clear that their fatty acid recognition is most likely defined by the size of their active site in combination with the availability of different fatty acids in the form of acyl-CoAs (Muszbek et al., 1999; Nuskova et al., 2021; Rana et al., 2019). It has been shown that some ZDHHC-PATs preferentially acylate with shorter fatty acids due to the presence of bulky amino acids in the cavity that accommodates their active site (Greaves et al., 2017; Jennings and Linder, 2012; Rana et al., 2018; Rana et al., 2019). Nonetheless, the target proteins we have studied in the past appear to be acylated redundantly by multiple ZDHHC-PATs, mitigating somewhat these effects.

To study protein *S*-acylation, we used an established protocol based on metabolic labeling with azido fatty acids and subsequent isolation of labeled proteins via click chemistry (**Figure 1A**). We previously showed that C16:0, C18:0 and C18:1, when provided in the culture medium, are all taken up by cells quite well, leading to changes in the endogenous intracellular levels of acyl-CoAs (Nuskova et al., 2021). Since exogenous palmitate increased C16:0-CoA levels but not C16:1-CoA levels, we assume that the modification with C15-az corresponds almost exclusively to palmitoylation. However, exogenous stearate increased C18:0-CoA and mainly C18:1-CoA. The modification with C17-az corresponds, therefore, more likely to oleoylation rather than to stearoylation. This is also supported by a finding that there is almost no stearoylation in HeLa and HEK293 cell lines, compared to palmitoylation or oleoylation (Schulte-Zweckel et al., 2019).

Of note, clickable fatty acids were used for large scale studies of *S*-acylated proteins in the past. Most studies employed 17-octadecyonic acid (17-ODYA) with 18 carbons (Martin and Cravatt, 2009) or 15-hexadecyonic acid (15-HDYA) that probably better mimics palmitate. Regardless of the length of the fatty acid analog, however, the studied modification was often called ‘palmitoylation’ (Suazo et al., 2021). In part since several studies reported that some ZDHHC-PATs do not transfer fatty acids longer than palmitate efficiently (Greaves et al., 2017; Jennings and Linder, 2012), it is probably safer to interpret the ‘palmitoylated’ proteins that result from these studies as ‘acylated’ proteins. To our knowledge, one study screened for proteins using 8-hour metabolic labeling with clickable fatty acids of different lengths in order to catch a larger set of acylated proteins (Wilson et al., 2011). Here, we focus specifically on differential acylation. We use 3-hour labeling to minimize the interconversion of different lipid species, and we study whether differential acylation can happen on single sites.

Our proteomic screen may include some false positives since we were not able to validate the acylation of one protein, STX5, using the APE assay (Figure 2Q). It is possible that such proteins are nonetheless acylated, but not on a cysteine residue. It is known that proteins can also be *N*-acylated (on lysine or N-terminal glycine) or *O*-acylated (on serine or threonine) (Chen et al., 2021). The acyl-switch APE method used for the validation of candidate proteins in Figure 2 is based on the hydrolysis of thioester bonds using hydroxylamine, hence it detects only *S*-acylation but not so much *N*- and *O*-acylation (Suazo et al., 2021). In contrast, the on-bead digestion used prior to mass spectrometry in our screen detects all acylated proteins. Hence it will be interesting to study in the future whether, STX5, for instance, is *N*-acylated or *O*-acylated.

We should note that our study tests the capacity of proteins to be acylated by particular fatty acid species when the fatty acids are presented to cells. The fact that two fatty acids can acylate a protein equally well does not mean that the stoichiometry of acylation in the cell with those two fatty acids is 1:1, since the stoichiometry also reflects the abundance of the two fatty acids *in vivo*. For instance, we find that GNAI3 can be comparably acylated by C15-az and C17-az yet *in vivo* we find by MS that GNAI3 is more abundantly palmitoylated than oleoylated (Nuskova et al., 2021). If we expose cells to C18:0, however, this ratio shifts towards oleoylation (Nuskova et al., 2021).

In sum, we find that most *S*-acylated proteins can be differentially acylated by C16 and C18 fatty acid species. This competitive differential acylation occurs on single cysteine sites on the proteins. Since the stoichiometry of acylation on a protein can shift depending on exposure of a cell to lipids in its environment (Nuskova et al., 2021; Senyilmaz-Tiebe et al., 2018), this raises the possibility that the function of a large number of proteins can be regulated by the lipid environment via this mechanism. Future work will be required to analyze whether differential *S*-acylation regulates the function of individual proteins.

## Methods

### Cell culture

HEK293E and Hela cells (CRL-10852 and CCL-2, respectively; ATCC), were cultivated in high-glucose DMEM (Gibco) supplemented with 10 % FCS (Sigma-Aldrich), 100 U/mL Penicillin and 100 μg/mL Streptomycin (Gibco) in a 37 °C incubator under stable 5 % CO_2_ atmosphere.

For SILAC experiments, HEK293E cells were cultured in DMEM without glutamine, lysine and arginine supplemented with 10 % dialyzed FCS, 100 U/mL Penicillin and 100 μg/mL Streptomycin, 1.737 mM proline, 2.5 mM glutamine and either unlabeled lysine (0.798 mM) and arginine (0.398 mM) (“light” medium), or ^2^H-labeled lysine and ^13^C-labeled arginine (“medium”), or ^13^C, ^15^N-labeled lysine and ^13^C, ^15^N-labeled arginine (“heavy”), purchased as a kit from Silantes. Before the click experiment was performed, labeling efficiency was checked by MS in cell lysates. For transfection of plasmids, Lipofectamine 2000 transfection reagent (Thermo Scientific) was used 24 h prior to an experiment. For use in cell culture, azido fatty acids were conjugated to BSA as in (Senyilmaz et al., 2015): Fatty acids were dissolved at a concentration of 50 mM in 100 mM NaOH at 95 °C. Once in solution, 100 μL of fatty acid were dropwise added onto 600 uL of 10 % fatty acid-free BSA pre-warmed to 50 °C. After mixing, the volume was filled up to 1 mL with water to reach the final fatty acid concentration of 5 mM. BSA-conjugated fatty acids were applied to cells at a concentration of 100 μM. Cell lines were tested negative for mycoplasma contamination (Eurofins Genomics, Germany). Cell lines were authenticated using Multiplex Cell Authentication by Multiplexion (Heidelberg, Germany) as described in (Castro et al., 2013).

### Detection of lipid modifications based on click chemistry

C15-az and C17-az modified proteins were pulled down using the Click Chemistry Capture Kit (Jena Bioscience) and alkyne agarose beads (Jena Bioscience). Cells were seeded in 10 cm culture dishes. For metabolic labeling, they were incubated for 3 h with 100 μM fatty acid analogs conjugated to BSA. The following fatty acid azides were used: C15-az (15-azidopentadecanoic acid) from Life Technologies and C17-az (17-azidoheptadecanoic acid) synthesized as in (Senyilmaz et al., 2015). Treatment with BSA was used as a negative control. After treatment, cells were briefly rinsed with PBS equilibrated to room temperature and lysed in 800 μL of urea-based lysis buffer supplemented with a protease inhibitor cocktail (Roche). Lysates were transferred into tubes, sonicated on ice for 3× 10 s (10 % amplitude) and cleared by centrifugation at 20,800 g for 5 min at 4 °C. The click reactions were assembled according to the protocol provided by the manufacturer, i.e. 800 μL lysate was mixed with 200 μL alkyne agarose beads and 1 mL reaction mixture. Samples were rotated end-over-end for 16–20 h at room temperature. Agarose beads were processed according to manufacturer’s instructions, including several washing steps, reduction, alkylation and on-bead trypsin digest. After the digest, eluates were transferred into new tubes and dried out.

Peptide pellets were dissolved in 8 M urea buffer containing Lys-C and trypsin for additional in-solution digestion and subsequently desalted using Sep-Pak cartridges. Peptides were loaded on a cartridge trap column, packed with Acclaim PepMap300 C18, 5 μm, 300 Å wide pore (Thermo Scientific) and separated via a gradient from 3% to 40% acetonitrile on a nanoEase MS Peptide analytical column (300 Å, 1.7 μm, 75 μm x 200 mm; Waters) using a 90 min MS-method. Eluting peptides were analyzed by an online-coupled Orbitrap Exploris 480 mass spectrometer.

Data analysis was carried out by MaxQuant (version 1.6.14.0). The data were filtered on an FDR cut-off of 0.01 on both peptide and protein level. In the conventional screen, quantification was done using a label free quantification (LFQ) approach based on the MaxLFQ algorithm. A minimum number of quantified peptides was required for protein quantification. For the SILAC analysis, match between runs option was enabled to transfer peptide identifications across Raw files based on accurate retention time and m/z. Quantification was done using a SILAC triplex approach with Arg6/Lys4 as medium and Arg10/Lys8 as heavy labeled amino acids. A minimum number of quantified peptides was required for protein quantification, the requantify option was activated to enable quantification of proteins with very high ratios.

To detect proteins pulled down on alkyne beads on a WB, the beads were washed with H_2_O after the Click reaction, pelleted at 1,000 g for 1 min and transferred into columns for extensive washing with the agarose washing buffer. Washed beads were resuspended in H_2_O and transferred into tubes again. Beads were pelleted at 1,000 g for 1 min and resuspended in 2× Laemmli sample buffer supplemented with 1 M hydroxylamine to elute bound proteins. Samples were incubated for 15 min at room temperature and then boiled at 95 °C for 5 min. Beads were pelleted at 1,000 g and the eluate was transferred into new tubes. Pulled-down proteins were loaded on a gel together with cell lysates that were kept as inputs. Western blots were probed with antibodies to test for the presence of specific proteins in pulldowns.

### Statistical analyses

To identify putative *S*-acylated proteins, we employed a mass spectrometry-based label-free strategy. Statistical analysis of the data was done using a free-access web application for quantitative analysis of proteomics data (https://proteomics.fgu.cas.cz/ProteoAnalyser/) (Villalba Sanchez et al., manuscript in preparation). For analysis, i*BAQ* values were used. The processing of the data followed removal of contaminants and a filtration step where the proteins that had more than 1 missing value in C15/C17-az labeled samples were filtered out, i.e. the data were filtered to keep at least 3 out of 4 valid values per protein in at least one of the labeled groups. Proteins that were exclusively found in C15/C17-az labeled samples, i.e. that had 5 missing values in the group of negative controls treated with BSA (n=5), were considered highly likely acylated and therefore included in the list of putative *S*-acylated proteins even if they were not significantly enriched according to the statistical analysis (the group of “clean” proteins). After the filtration, data were log-transformed (base 2), normalized by variance stabilizing transformation (VSN) and missing values were imputed using the minimum value method. To perform Bayesian statistical tests, the *limma* package was used. Proteins significantly enriched in C15/C17-az labeled samples over negative controls (p_adj_<0.05) were considered putative *S*-acylated proteins.

### SILAC analysis

Analysis was conducted on the SILAC data after pre-processing by the MaxQuant software with no further normalization nor filtering. The normalized ratios were first log-transformed (base 2), afterwards, ratios with flipped denominators and numerators were multiplied by -1 to ensure the comparability of all ratios. Finally, the analysis was conducted in two-parts; the first stage consisted of a global test comparing C15/C17-az labeled samples with negative controls. In the second stage, proteins that were found to be significantly enriched in C15-az and/or C17-az labeled samples compared negative controls (padj<0.05) were further analyzed to compare C15-az and C17-az labeled samples using the unadjusted p-values. In each stage, the limma package was used to estimate the fold changes and standard errors by fitting a linear model for each protein. The method for two-stage p-value adjustment was selected using a simulation study to compare the effect of multiple adjustment strategies on the false discovery rate and the proportion of missed hits.

### Cloning and site-directed mutagenesis

Sequences of primers used in this study are provided in **Supplementary Table 3**. The open reading frames of human *HRAS*, *LAMTOR1*, and *TFR1* were PCR amplified from cDNA of HeLa cells with primers containing EcoRI and NotI (*HRAS*), XhoI and BglII (*LAMTOR1*) and ClaI and BglII (*TFR1*) restriction sites using Phusion High-Fidelity Polymerase (Thermo Scientific) and cloned into a pcDNA3 backbone containing an N-terminal V5-tag (*HRAS*) or C-terminal HA-tag (*LAMTOR1, TFR1*). To mutate the LAMTOR1 acylation sites, primers with the desired mutation were used to PCR amplify *LAMTOR1* from pcDNA3-*LAMTOR1-HA* and the amplified mutated inserts were cloned into the same pcDNA3 backbone containing a C-terminal HA-tag. To mutate the acylation sites of *HRAS*, primers with the desired mutations were used to PCR amplify the whole construct containing plasmids. Site-directed mutagenesis on *TFR1* was performed by overlap extension PCR. The sequence of all constructs was confirmed by sequencing (Microsynth, Germany). Primers were designed using the primer design tool from Takara Bio (https://www.takarabio.com/learning-centers/cloning/primer-design-and-other-tools). Sequence manipulation was performed using A plasmid Editor (ApE) software.

### Acyl-PEG exchange (APE) assay

To detect *S*-acylation, the acyl-PEG exchange assay, a method based on selective maleimide-labeling of acylated cysteines causing a mass shift, was performed as described in (Percher et al., 2016; Percher et al., 2017), using methoxypolyethylene glycol maleimide (mPEG, 5 kDa).

### Protein electrophoresis and Western blotting

Proteins were separated with Tris-glycine SDS-PAGE, transferred onto a nitrocellulose membrane and detected with primary antibodies according to manufacturers’ instructions followed by incubation with HRP-conjugated secondary antibodies diluted 1:10,000 (Jackson ImmunoResearch Labs). Chemiluminescence was recorded with the Chemidoc Imager (Bio-Rad) and quantified using Image Lab software (Bio-Rad). Rabbit polyclonal anti-CYB5B (#15469-1-AP), anti-DR4 (#24063-1-AP), anti-FAS/CD95 (#13098-1-AP), anti-HRAS (#18295-1-AP), anti-AEG-1/MTDH (#13860-1-AP), anti-RFFL (#12687-1-AP), anti-RRAS2 (#12530-1-AP), anti-SCAMP3 (#26888-1-AP), anti-STX12 (#14259-1-AP), anti-STX5 (#26711-1-AP), anti-STX7 (#12322-1-AP), anti-TMX1 (#27489-1-AP), anti-VAMP3/Cellubrevin (#10702-1-AP), anti-VAMP4 (#10738-1-AP) antibodies were purchased from Proteintech. Rabbit monoclonal anti-ASCT2 (SLC1A5; #5245), anti-CD71 (Transferrin receptor 1; #13113), anti-HA tag (#3724S), and anti-LAMTOR1 (#8975) antibodies were purchased from Cell Signaling Technology. Rabbit monoclonal anti-GNAI2 (#ab137050) and mouse monoclonal anti-calnexin (ab31290) antibodies were purchased from Abcam. Mouse monoclonal anti-V5 tag (#R960-25), rabbit monoclonal anti-LIMP2 (SCARB2; #702770) and rabbit polyclonal anti-FASN (#PA5-22061) antibodies were purchased from ThermoFisher. Mouse monoclonal anti-IRS4 (#sc-100854) antibody was purchased from Santa Cruz, mouse monoclonal anti-α-tubulin (#T9026, 1:3,000) from Sigma-Aldrich, and rabbit polyclonal anti-NPC1 (#NB400-148SS) from Novus Biologicals. All primary antibodies were diluted 1:1,000 unless stated differently.

### Figure preparation

Figures were prepared using Affinity Designer (https://affinity.serif.com/en-gb/).

## ACKNOWLEDGMENTS

This project was funded by the European Research Council (ERC) under the European Union’s Horizon 2020 research and innovation program (grant agreement No 724286 - C18Signaling) to AAT.

## AUTHOR CONTRIBUTIONS

H.N., F.G.C., L.S.S., M.T., A.K.M., A.A.T. designed experiments, H.N., F.G.C., L.S.S., M.T., M.S., performed experiments, H.N., F.G.C., L.S.S., M.T., C.R., M.S., D.H., and A.A.T. analyzed data, H.N. and A.A.T wrote the manuscript.

## DATA AVAILABILITY

All data generated in this study are provided in the Supplementary Information and Source Data files.

## Abbreviations

ABE: acyl-biotin exchange
APE: acyl-PEG exchange
APTs: acyl protein thioesterases
MS: mass spectrometry
PPTs: palmitoyl protein thioesterases
RAC: resin-assisted capture
TFR1: transferrin receptor 1
ZDHHC-PATs: zinc finger aspartate-histidine-histidine-cysteine (DHHC) domain containing family of protein acyl transferases

## REFERENCES

Ahearn, I., Zhou, M., and Philips, M.R. 2018. Posttranslational Modifications of RAS Proteins. Cold Spring Harb Perspect Med 8. 10.1101/cshperspect.a031484

Alvarez, E., Girones, N., and Davis, R.J. 1990. Inhibition of the receptor-mediated endocytosis of diferric transferrin is associated with the covalent modification of the transferrin receptor with palmitic acid. The Journal of biological chemistry 265: 16644–55.

Blanc, M., David, F.P.A., and van der Goot, F.G. 2019. SwissPalm 2: Protein S-Palmitoylation Database. Methods Mol Biol 2009: 203–14. 10.1007/978-1-4939-9532-5_16

Busquets-Hernandez, C., and Triola, G. 2021. Palmitoylation as a Key Regulator of Ras Localization and Function. Front Mol Biosci 8: 659861. 10.3389/fmolb.2021.659861

Castro, F., Dirks, W.G., Fahnrich, S., Hotz-Wagenblatt, A., Pawlita, M., and Schmitt, M. 2013. High-throughput SNP-based authentication of human cell lines. Int J Cancer 132: 308–14. 10.1002/ijc.27675

Chamberlain, L.H., and Shipston, M.J. 2015. The physiology of protein S-acylation. Physiological reviews 95: 341–76. 10.1152/physrev.00032.2014

Charron, G., Zhang, M.M., Yount, J.S., Wilson, J., Raghavan, A.S., Shamir, E., and Hang, H.C. 2009. Robust fluorescent detection of protein fatty-acylation with chemical reporters. J Am Chem Soc 131: 4967–75. 10.1021/ja810122f

Chen, B., Sun, Y., Niu, J., Jarugumilli, G.K., and Wu, X. 2018. Protein Lipidation in Cell Signaling and Diseases: Function, Regulation, and Therapeutic Opportunities. Cell Chem Biol 25: 817–31. 10.1016/j.chembiol.2018.05.003

Chen, J.J., Fan, Y., and Boehning, D. 2021. Regulation of Dynamic Protein S-Acylation. Front Mol Biosci 8: 656440. 10.3389/fmolb.2021.656440

Ciechanover, A., Schwartz, A.L., and Lodish, H.F. 1983. Sorting and recycling of cell surface receptors and endocytosed ligands: the asialoglycoprotein and transferrin receptors. J Cell Biochem 23: 107–30. 10.1002/jcb.240230111

Degtyarev, M.Y., Spiegel, A.M., and Jones, T.L. 1993. Increased palmitoylation of the Gs protein alpha subunit after activation by the beta-adrenergic receptor or cholera toxin. J Biol Chem 268: 23769–72.

Dixon, C.L., Mekhail, K., and Fairn, G.D. 2021. Examining the Underappreciated Role of S-Acylated Proteins as Critical Regulators of Phagocytosis and Phagosome Maturation in Macrophages. Front Immunol 12: 659533. 10.3389/fimmu.2021.659533

Drisdel, R.C., and Green, W.N. 2004. Labeling and quantifying sites of protein palmitoylation. Biotechniques 36: 276–85. 10.2144/04362RR02

El-Husseini Ael, D., Schnell, E., Dakoji, S., Sweeney, N., Zhou, Q., Prange, O., Gauthier-Campbell, C., Aguilera-Moreno, A., Nicoll, R.A., and Bredt, D.S. 2002. Synaptic strength regulated by palmitate cycling on PSD-95. Cell 108: 849–63. 10.1016/s0092-8674(02)00683-9

Essandoh, K., Philippe, J.M., Jenkins, P.M., and Brody, M.J. 2020. Palmitoylation: A Fatty Regulator of Myocardial Electrophysiology. Front Physiol 11: 108. 10.3389/fphys.2020.00108

Figlia, G., Willnow, P., and Teleman, A.A. 2020. Metabolites Regulate Cell Signaling and Growth via Covalent Modification of Proteins. Developmental cell 54: 156–70. 10.1016/j.devcel.2020.06.036

Fritsch, J., Sarchen, V., and Schneider-Brachert, W. 2021. Regulation of Death Receptor Signaling by S-Palmitoylation and Detergent-Resistant Membrane Micro Domains-Greasing the Gears of Extrinsic Cell Death Induction, Survival, and Inflammation. Cancers (Basel) 13. 10.3390/cancers13112513

Fujimoto, T., Stroud, E., Whatley, R.E., Prescott, S.M., Muszbek, L., Laposata, M., and McEver, R.P. 1993. P-selectin is acylated with palmitic acid and stearic acid at cysteine 766 through a thioester linkage. The Journal of biological chemistry 268: 11394–400.

Greaves, J., Munro, K.R., Davidson, S.C., Riviere, M., Wojno, J., Smith, T.K., Tomkinson, N.C., and Chamberlain, L.H. 2017. Molecular basis of fatty acid selectivity in the zDHHC family of S-acyltransferases revealed by click chemistry. Proceedings of the National Academy of Sciences of the United States of America 114: E1365–E74. 10.1073/pnas.1612254114

Hallak, H., Muszbek, L., Laposata, M., Belmonte, E., Brass, L.F., and Manning, D.R. 1994. Covalent binding of arachidonate to G protein alpha subunits of human platelets. J Biol Chem 269: 4713–6.

Han, J., Pluhackova, K., and Bockmann, R.A. 2017. The Multifaceted Role of SNARE Proteins in Membrane Fusion. Front Physiol 8: 5. 10.3389/fphys.2017.00005

Hancock, J.F., Magee, A.I., Childs, J.E., and Marshall, C.J. 1989. All ras proteins are polyisoprenylated but only some are palmitoylated. Cell 57: 1167–77. 10.1016/0092-8674(89)90054-8

Hang, H.C., Geutjes, E.J., Grotenbreg, G., Pollington, A.M., Bijlmakers, M.J., and Ploegh, H.L. 2007. Chemical probes for the rapid detection of Fatty-acylated proteins in Mammalian cells. J Am Chem Soc 129: 2744–5. 10.1021/ja0685001

He, Y., and Linder, M.E. 2009. Differential palmitoylation of the endosomal SNAREs syntaxin 7 and syntaxin 8. J Lipid Res 50: 398–404. 10.1194/jlr.M800360-JLR200

Jennings, B.C., and Linder, M.E. 2012. DHHC protein S-acyltransferases use similar ping-pong kinetic mechanisms but display different acyl-CoA specificities. The Journal of biological chemistry 287: 7236–45. 10.1074/jbc.M111.337246

Ko, P.J., and Dixon, S.J. 2018. Protein palmitoylation and cancer. EMBO Rep 19. 10.15252/embr.201846666

Liang, X., Lu, Y., Neubert, T.A., and Resh, M.D. 2002. Mass spectrometric analysis of GAP-43/neuromodulin reveals the presence of a variety of fatty acylated species. J Biol Chem 277: 33032–40. 10.1074/jbc.M204607200

Liang, X., Nazarian, A., Erdjument-Bromage, H., Bornmann, W., Tempst, P., and Resh, M.D. 2001. Heterogeneous fatty acylation of Src family kinases with polyunsaturated fatty acids regulates raft localization and signal transduction. The Journal of biological chemistry 276: 30987–94. 10.1074/jbc.M104018200

Lin, D.T., and Conibear, E. 2015. ABHD17 proteins are novel protein depalmitoylases that regulate N-Ras palmitate turnover and subcellular localization. Elife 4: e11306. 10.7554/eLife.11306

Martin, B.R., and Cravatt, B.F. 2009. Large-scale profiling of protein palmitoylation in mammalian cells. Nat Methods 6: 135–8. 10.1038/nmeth.1293

Montigny, C., Decottignies, P., Le Marechal, P., Capy, P., Bublitz, M., Olesen, C., Moller, J.V., Nissen, P., and le Maire, M. 2014. S-palmitoylation and s-oleoylation of rabbit and pig sarcolipin. J Biol Chem 289: 33850–61. 10.1074/jbc.M114.590307

Muszbek, L., Haramura, G., Cluette-Brown, J.E., Van Cott, E.M., and Laposata, M. 1999. The pool of fatty acids covalently bound to platelet proteins by thioester linkages can be altered by exogenously supplied fatty acids. Lipids 34 **Suppl**: S331–7. 10.1007/bf02562334

Nada, S., Hondo, A., Kasai, A., Koike, M., Saito, K., Uchiyama, Y., and Okada, M. 2009. The novel lipid raft adaptor p18 controls endosome dynamics by anchoring the MEK-ERK pathway to late endosomes. EMBO J 28: 477–89. 10.1038/emboj.2008.308

Nuskova, H., Serebryakova, M.V., Ferrer-Caelles, A., Sachsenheimer, T., Luchtenborg, C., Miller, A.K., Brugger, B., Kordyukova, L.V., and Teleman, A.A. 2021. Stearic acid blunts growth-factor signaling via oleoylation of GNAI proteins. Nature communications 12: 4590. 10.1038/s41467-021-24844-9

Percher, A., Ramakrishnan, S., Thinon, E., Yuan, X., Yount, J.S., and Hang, H.C. 2016. Mass-tag labeling reveals site-specific and endogenous levels of protein S-fatty acylation. Proceedings of the National Academy of Sciences of the United States of America 113: 4302–7. 10.1073/pnas.1602244113

Percher, A., Thinon, E., and Hang, H. 2017. Mass-Tag Labeling Using Acyl-PEG Exchange for the Determination of Endogenous Protein S-Fatty Acylation. Curr Protoc Protein Sci 89: 14 17 1–14 17 11. 10.1002/cpps.36

Rana, M.S., Kumar, P., Lee, C.J., Verardi, R., Rajashankar, K.R., and Banerjee, A. 2018. Fatty acyl recognition and transfer by an integral membrane S-acyltransferase. Science 359. 10.1126/science.aao6326

Rana, M.S., Lee, C.J., and Banerjee, A. 2019. The molecular mechanism of DHHC protein acyltransferases. Biochem Soc Trans 47: 157–67. 10.1042/BST20180429

Rocks, O., Gerauer, M., Vartak, N., Koch, S., Huang, Z.P., Pechlivanis, M., Kuhlmann, J., Brunsveld, L., Chandra, A., Ellinger, B., et al. 2010. The palmitoylation machinery is a spatially organizing system for peripheral membrane proteins. Cell 141: 458–71. 10.1016/j.cell.2010.04.007

Roth, A.F., Wan, J., Green, W.N., Yates, J.R., and Davis, N.G. 2006. Proteomic identification of palmitoylated proteins. Methods 40: 135–42. 10.1016/j.ymeth.2006.05.026

Sancak, Y., Bar-Peled, L., Zoncu, R., Markhard, A.L., Nada, S., and Sabatini, D.M. 2010. Ragulator-Rag complex targets mTORC1 to the lysosomal surface and is necessary for its activation by amino acids. Cell 141: 290–303. 10.1016/j.cell.2010.02.024

Sanders, S.S., De Simone, F.I., and Thomas, G.M. 2019. mTORC1 Signaling Is Palmitoylation-Dependent in Hippocampal Neurons and Non-neuronal Cells and Involves Dynamic Palmitoylation of LAMTOR1 and mTOR. Front Cell Neurosci 13: 115. 10.3389/fncel.2019.00115

Schulte-Zweckel, J., Dwivedi, M., Brockmeyer, A., Janning, P., Winter, R., and Triola, G. 2019. A hydroxylamine probe for profiling S-acylated fatty acids on proteins. Chem Commun (Camb) 55: 11183–86. 10.1039/c9cc05989j

Segal-Salto, M., Sapir, T., and Reiner, O. 2016. Reversible Cysteine Acylation Regulates the Activity of Human Palmitoyl-Protein Thioesterase 1 (PPT1). PLoS One 11: e0146466. 10.1371/journal.pone.0146466

Senyilmaz, D., Virtue, S., Xu, X., Tan, C.Y., Griffin, J.L., Miller, A.K., Vidal-Puig, A., and Teleman, A.A. 2015. Regulation of mitochondrial morphology and function by stearoylation of TFR1. Nature 525: 124–8. 10.1038/nature14601

Senyilmaz-Tiebe, D., Pfaff, D.H., Virtue, S., Schwarz, K.V., Fleming, T., Altamura, S., Muckenthaler, M.U., Okun, J.G., Vidal-Puig, A., Nawroth, P., et al. 2018. Dietary stearic acid regulates mitochondria in vivo in humans. Nature communications 9: 3129. 10.1038/s41467-018-05614-6

Suazo, K.F., Park, K.Y., and Distefano, M.D. 2021. A Not-So-Ancient Grease History: Click Chemistry and Protein Lipid Modifications. Chem Rev 121: 7178–248. 10.1021/acs.chemrev.0c01108

Wan, J., Roth, A.F., Bailey, A.O., and Davis, N.G. 2007. Palmitoylated proteins: purification and identification. Nature protocols 2: 1573–84. 10.1038/nprot.2007.225

Wilson, J.P., Raghavan, A.S., Yang, Y.Y., Charron, G., and Hang, H.C. 2011. Proteomic analysis of fatty-acylated proteins in mammalian cells with chemical reporters reveals S-acylation of histone H3 variants. Mol Cell Proteomics 10: M110 001198. 10.1074/mcp.M110.001198

Won, S.J., Cheung See Kit, M., and Martin, B.R. 2018. Protein depalmitoylases. Crit Rev Biochem Mol Biol 53: 83–98. 10.1080/10409238.2017.1409191

Zareba-Koziol, M., Figiel, I., Bartkowiak-Kaczmarek, A., and Wlodarczyk, J. 2018. Insights Into Protein S-Palmitoylation in Synaptic Plasticity and Neurological Disorders: Potential and Limitations of Methods for Detection and Analysis. Front Mol Neurosci 11: 175. 10.3389/fnmol.2018.00175

Zhang, M.M., and Hang, H.C. 2017. Protein S-palmitoylation in cellular differentiation. Biochem Soc Trans 45: 275–85. 10.1042/BST20160236

Zhang, M.M., Tsou, L.K., Charron, G., Raghavan, A.S., and Hang, H.C. 2010. Tandem fluorescence imaging of dynamic S-acylation and protein turnover. Proc Natl Acad Sci U S A 107: 8627–32. 10.1073/pnas.0912306107

Zheng, B., Jarugumilli, G.K., Chen, B., and Wu, X. 2016. Chemical Probes to Directly Profile Palmitoleoylation of Proteins. Chembiochem 17: 2022–27. 10.1002/cbic.201600403

